# Chronic β3 adrenergic agonist treatment improves brain microvascular endothelial function and cognition in aged mice

**DOI:** 10.1101/2024.07.09.602747

**Authors:** Duraipandy Natarajan, Shoba Ekambaram, Stefano Tarantini, Raghavendra Yelahanka Nagaraja, Andriy Yabluchanskiy, Andria F Hedrick, Vibhudutta Awasthi, Madhan Subramanian, Anna Csiszar, Priya Balasubramanian

**Author notes:** Correspondence: Dr. Priya Balasubramanian Assistant Professor Department of Neurosurgery Oklahoma Center for Geroscience and Healthy Brain Aging University of Oklahoma Health Sciences Center 940 Stanton L. Young Blvd, BMSB 842 Oklahoma City, OK 73104.

## Abstract

Microvascular endothelial dysfunction, characterized by impaired neurovascular coupling, reduced glucose uptake, blood-brain barrier disruption, and microvascular rarefaction, plays a critical role in the pathogenesis of age-related vascular cognitive impairment (VCI). Emerging evidence points to non-cell autonomous mechanisms mediated by adverse circulating milieu (an increased ratio of pro-geronic to anti-geronic circulating factors) in the pathogenesis of endothelial dysfunction leading to impaired cerebral blood flow and cognitive decline in the aging population. In particular, age-related adipose dysfunction contributes, at least in part, to an unfavorable systemic milieu characterized by chronic hyperglycemia, hyperinsulinemia, dyslipidemia, and altered adipokine profile, which together contribute to microvascular endothelial dysfunction. Hence, in the present study, we aimed to test whether thermogenic stimulation, an intervention known to improve adipose and systemic metabolism by increasing cellular energy expenditure, could mitigate brain endothelial dysfunction and improve cognition in the aging population. Eighteen-month-old old C57BL/6J mice were treated with saline or CL (β3-adrenergic agonist) for 6 weeks followed by functional analysis to assess endothelial function and cognition. CL treatment improved neurovascular coupling responses and rescued brain glucose uptake in aged animals. In addition, CL treatment also attenuated blood-brain barrier leakage and associated neuroinflammation in the cortex of aged animals. More importantly, these beneficial changes in microvascular function translated to improved cognitive performance in radial arm water maze and Y-maze tests. Our results suggest that β3-adrenergic agonist treatment improves multiple aspects of brain microvascular endothelial function and can be potentially repurposed for treating age-associated cognitive decline.

## Introduction

Microvascular endothelial cells, which form the inner lining of all cerebral vessels, play multifaceted roles in regulating cerebral blood flow and cognition. First, endothelial nitric oxide contributes to neurovascular coupling responses (NVC), a critical vasodilatory mechanism that maintains neuronal homeostasis and function by promptly matching local neuronal activity with the required increase in cerebral blood flow (CBF) ^1,2^. Secondly, endothelial glucose uptake through Glut1 (endothelial isoform 55kDa) controls the first point of glucose entry into the brain and critically contributes to the maintenance of whole-brain energy homeostasis in addition to supporting its metabolic needs^3^. Thirdly, endothelial cells maintain the structural integrity of the blood-brain barrier (BBB), which prevents the entry of serum constituents into the brain parenchyma, and subsequent glial activation and neuroinflammation^4,5^. Lastly, endothelial angiogenesis is key to maintaining the optimal cerebral microvascular density needed to achieve adequate perfusion to the entire brain^6^. Accumulating evidence points to age-related impairment in endothelial function, metabolism, and structure resulting in impaired NVC^7,8^, attenuated glucose uptake leading to hypometabolism^9,10^, BBB leakage leading to neuroinflammation^11,12^ and microvascular rarefaction^13,14^. All these mechanisms synergistically act to reduce cerebral blood flow or perfusion and contribute to cognitive decline in aging.

The majority of the previous research in the microvascular aging field has primarily focused on targeting cell-intrinsic mechanisms including senescence, oxidative stress, DNA damage, etc. However, a paradigm shift in the mechanistic view of endothelial dysfunction has occurred since the emergence of results from heterochronic parabiosis studies in recent years. These studies where young and old mice share the circulation for an extended period highlight the pro-aging role of circulating factors in accelerating endothelial dysfunction^15–19^. Especially, altered systemic and metabolic milieu in aging including chronic inflammation, hyperglycemia, hyperinsulinemia, and dyslipidemia has been implicated in endothelial dysfunction^20–25^. In addition, age-related reduction in the circulating vasoprotective factors like IGF1 and adiponectin could also potentially contribute to microvascular aging^26–28^. Conforming to this overall idea, several meta-analysis studies have reported that patients diagnosed with age-related diabetes, dyslipidemia, and metabolic syndrome have an elevated risk of developing cognitive impairment later in their lifetime^29–32^. Overall, these studies signify the importance of interventions that improve metabolic dysfunction in treating and/or preventing VCI in aging.

Adipose tissue plays a central role in whole-body energy homeostasis through its direct involvement in glucose and lipid metabolism and also indirectly via its crosstalk with other systemic tissues through secreted factors. Age-related pathological changes in adipose tissue contribute to metabolic dysfunction through ectopic lipid deposition, insulin resistance, and low-grade chronic inflammation^28,33–36^, all of which have been implicated in accelerating endothelial aging. On the other hand, improvements in adipose tissue metabolism, at least in part, contribute to the delayed aging phenotype observed in response to several well-known anti-aging interventions^28,36–38^. More importantly, adipose-related metabolic dysfunction in middle age precedes the onset of cognitive decline later in life^39–42^, suggesting that interventions that restore adipose and systemic metabolism could be targeted to delay or prevent brain endothelial dysfunction and VCI in aging.

We have recently reported that the pharmacological activation of adipose thermogenesis, a catabolic phenomenon marked by increased fuel oxidation and energy expenditure, improved the overall systemic metabolism in aged mice^43^. Specifically, we used a β3-adrenergic receptor agonist (β3AR, CL, 216243) to stimulate thermogenesis in aged mice. β3ARs are predominantly expressed in the white and brown adipose tissue and play a critical role in the maintenance and activation of lipolytic and thermogenic machinery. β3AR agonists (CL, 316243 in rodents and FDA-approved mirabegron in humans) have been extensively validated as a pharmacological means to stimulate thermogenesis and improve systemic glucose and lipid metabolism^44–46^, however, its relevance in aged subjects has remained questionable. Addressing this, we have recently shown that the metabolic benefits of β3AR agonist treatment are preserved in aged mice. Chronic β3AR stimulation increased whole-body energy expenditure, reduced fat mass, improved glucose tolerance and insulin sensitivity, increased circulating adiponectin levels, and reduced ectopic lipid deposition in aged mice^43^. In the present study, we wanted to examine whether β3AR stimulation-mediated improvements in systemic metabolism and circulating milieu can mitigate microvascular endothelial dysfunction and cognitive decline in aged mice.

## Materials and Methods

### Animals and treatment

All animal protocols were approved by the Institutional Animal Care and Use Committee at the University of Oklahoma Health Sciences Center (OUHSC). Aged C57BL/6JN male mice (18 months old) were obtained from the aging colony maintained by the National Institute on Aging at Charles River Laboratories (Wilmington, MA) and were fed a standard chow diet (PicoLab Rodent Diet 5053) with continuous access to water and enrichment. The animals were housed in the conventional animal housing facility with a 12:12-hour light-dark cycle at OUHSC. Aged mice were implanted with osmotic minipumps filled with saline or β3-AR agonist [CL 316,243 (CL) R&D Systems-Cat. No. 1499/50, 0.75 nmol/h] to enable continuous infusion for 6 weeks as previously described^46^. At the end of 4 weeks of treatment, the mice were subjected to behavioral assays including radial arm water maze (RAWM) and Y-maze to assess spatial learning and memory-related cognitive outcomes. A sub-cohort of animals underwent PET/CT imaging to assess glucose uptake at the end of 4 weeks. All the animals were sacrificed at the end of 6 weeks and brain tissues were collected and either stored at -80C for protein analysis or fixed in 10% formalin for paraffin embedding. A sepatate cohort of young (3-4 mos) and aged (20-22 mos) C57BL/6J animals were also used for validation of PET/CT imaging technique.

### Radial arm water maze test

Spatial memory and long-term memory in mice were assessed by performance in the radial arm water maze (RAWM) test as described previously^47^. The RAWM consisted of eight 9 cm wide arms that radiated out from the open central area, with a submerged escape platform located at the end of one of the arms. Food-grade white paint was added to make the water opaque and mask the escape platform. Visual cues were marked inside the maze at the end of each arm. The movement of mice was monitored by a video tracking system directly above the maze and the parameters including distance, time, and latency to escape were recorded using Ethovision software (Noldus Information Technology Inc., Leesburg, VA, USA). The experiment comprised 3 consecutive days of learning trials, followed by a 1-week break, then 1 day of probe trial (recalling), and the last day of relearning (reversal). During the learning period, mice were allowed to learn the location of the submerged platform during two session blocks each consisting of four consecutive acquisition trials. On each trial, the mouse started in one arm not containing the platform, and was allowed to wade for up to 1 minute to find the escape platform. All mice were trained to identify the platform following 1st trial of day 1. During the probe trial, the mice were subjected to test their memory by recalling the learning trial. On the next day (reversal trial), the ability of the mice to relearn the task, when the platform had been moved to a different arm that was not adjacent nor diametrically positioned to the previous location was assessed. The mice were charged an error whenever they entered an incorrect arm (all four paws within the distal half of the arm) or spent 15 seconds at the center without entering any arm.

### Y maze test

Mice were also assessed for spatial reference memory using a Y maze test as described previously^48^. The Y-maze apparatus consisted of three arms that were positioned 120 degrees apart. The walls of the arms were 15 cm high, allowing the mouse to see distal spatial landmarks. The test consisted of a learning trial and a test trial separated by 4 hours. During the learning trial, one of the arms was closed off and the mice were allowed to freely explore the other 2 arms for 5 minutes. After 4 hours, the obstruction to one of the arms was removed and the mice were allowed to explore all 3 arms for 5 minutes (test trial). The arm that was obstructed during the learning trial is the novel arm and the other 2 arms are the familiar arms. The mouse should remember the unexplored novel arm during the test trial and spend more time exploring this arm (spatial reference memory). The amount of time spent in the novel arm and the number of novel arm entries were calculated from the video recording using the Ethovision software (Noldus Information Technology Inc., Leesburg, VA, USA).

### Neurovascular coupling assessments

After completion of behavioral tests, neurovascular coupling responses were assessed in a sub-group of animals using laser speckle contrast imaging as described previously^48,49^. Briefly, mice were anesthetized, endotracheally intubated and ventilated. Vitals including core body temperature, blood pressure and blood gas were monitored throughout the experiment to ensure that they remain within the physiological range. Following immobilization in the stereotaxic frame, the scalp and periosteum were opened and the skull was thinned using a dental drill. To avoid overheating during drilling, dripping buffer was infused at the drilling site. After placement of the laser speckle contrast imager (Perimed, Jarfalla, Sweden) above the thinned site, CBF responses on the left somatosensory cortex were captured by stimulating the right whiskers for 30s at 5Hz from side to side. A total of six trials were performed with 5-10-minute intervals between them. The average of the CBF changes during the 6 trials was expressed as a % increase from the baseline values.

### ^18^F-FDG PET/CT imaging to assess brain glucose uptake

Briefly, overnight-fasted animals were injected with ^18^F-FDG (100 μCi) via the tail vein. After 2 hours of FDG uptake, a 15-minute PET image was acquired immediately followed by a 2-minute CT image. Both images were acquired using an MI Labs Vector6 machine (Utrecht, Netherlands). Images were reconstructed and registered using MI Labs software. ROI for the brain was then manually selected and the ^18^F activity in this ROI was quantified in the corresponding region of the registered PET image using AMIRA software (Thermofisher Scientific). Percent injected dose (% ID) was calculated as the activity (µCi) in the BAT decay corrected to the time of the injection (i.t.) divided by the injected activity (µCi). The standard uptake value (SUV) was calculated by normalizing the % ID to the body weight of the animals. The imaging and analysis were performed in the rodent imaging facility (RIF) at the College of Pharmacy in OUHSC.

### BBB permeability assays

BBB permeability was assessed by quantifying the levels of extravasated fluorescent tracers in brain lysates as described previously by Devraj et al^50^. Briefly, the mice were injected with 100µl of 2mM of 3Kda FITC dextran tracer (#D3305, Thermoscientific, Waltham, MA) by intraperitoneal injection. After 15 minutes, the mice were anesthetized and cardiac perfusion with ice-cold PBS was performed to remove the tracers from the vascular compartment. Cortex and hippocampus regions of the brain were dissected from one sagittal section of hemibrain and the other half of the brain was stored in formalin for immunofluorescence analysis. The permeability index was assessed by measuring the fluorescence intensity in the serum and brain homogenates (cortex and hippocampus) at an excitation/emission (nm) value of 490/520 using a plate reader. All raw fluorescence values (RFU) were corrected for background using tissue homogenates or serum from sham animals that did not receive tracer injection. The permeability index was calculated using the following formula: Permeability Index (mL/g) = (Tissue RFUs/g tissue weight)/ (Serum RFUs/mL serum).

### Capillary-based immunoassay for Glut 1 protein expression

Frozen cortex and hippocampus samples were lysed using 1X cell lysis buffer (Cell Signaling Technology) containing Halt protease and phosphatase inhibitor cocktail (Thermo Scientific, #PI78440). The lysates were obtained by mincing the tissue using a Dounce homogenizer followed by centrifugation at 16,000 g for 10 min at 4 °C. The clear supernatant was collected and the protein concentrations were determined using the Pierce BCA Protein Assay Kit (Thermo Scientific, #23227). Automated western blots were performed using Jess capillary-based immunoassay using 12-230 kDa separation with protein normalization (PN) module using the Compass for SW Software 6.2.0 (Protein Simple). Protein samples were diluted with 0.1X sample buffer and loaded at 0.5 mg/mL optimized concentration. Anti-GLUT1 antibody (Abcam #ab115730) was loaded at 1:50 dilution. The peak area for 45kDa and 55kDa isoforms of GLUT1 was calculated after normalization to the total protein content using the dropped line peak integration method available in the Compass for SW Software 6.2.0 (Protein Simple).

### Milliplex assays for cytokine analysis

Protein lysates from cortex samples were analyzed for inflammatory markers (Millipore Sigma #MCYTOMAG-70K-PMX) using Milliplex kits. The values from protein lysates were normalized to the total protein content in each sample assessed by BCA assay and expressed as pg/ug of protein.

### Statistical analysis

Statistical analyses were performed using Graph pad prism 9.3.1 (GraphPad Software, San Diego, CA, USA) and the data are expressed as mean±SEM. Data were analyzed by two-tailed, unpaired student’s t-test and p<0.05 were considered statistically significant.

## Results

### Chronic β3AR stimulation improved neurovascular coupling responses and brain glucose uptake in aged mice

To determine the impact of metabolic improvements on brain microvascular endothelial function, we assessed neurovascular coupling in the somatosensory cortex via laser speckle contrast imaging following 6 weeks of CL treatment. Cerebral blood flow responses in the somatosensory cortex in response to contralateral whisker stimulation were significantly increased in aged mice treated with CL when compared with age-matched controls (representative pseudocolor flowmetry maps are shown in Fig. 1A and the summary data are shown in Fig. 1B). Next, we utilized ^18^F-FDG PET/CT imaging to measure *in vivo* brain glucose uptake, another critical endothelial function mediated by glucose transporters expressed on the luminal surface that regulates whole-brain energy metabolism. First, we validated the ^18^F-FDG PET/CT imaging technique to detect age-related decreases in brain glucose uptake. In agreement with previous studies^51^, we were able to demonstrate a significant decrease in brain glucose uptake in aged animals compared with young controls (Suppl. Fig. 1A). Following validation of the imaging technique, we investigated whether CL treatment improved brain glucose uptake in aged animals. Consistent with improved NVC, brain glucose uptake was also significantly improved in aged animals following CL treatment (representative PET/CT images are shown in Fig. 1C and the summary data are shown in Fig. 1D). Glucose uptake at the BBB is mediated by GLUT1 transporter, which has 2 isoforms: 55kDa isoform expressed in the luminal side of the BBB endothelial cells and the 45kDa isoform expressed in the astrocytic end-feet. Correlating with increased brain glucose uptake, CL treatment significantly increased GLUT1 levels in the hippocampus of the aged mice. However, interestingly only the 55kDa endothelial GLUT1 isoform was upregulated while no changes were observed with the 45kDa astrocytic isoform (Fig. 1E-G).

**Figure 1:**
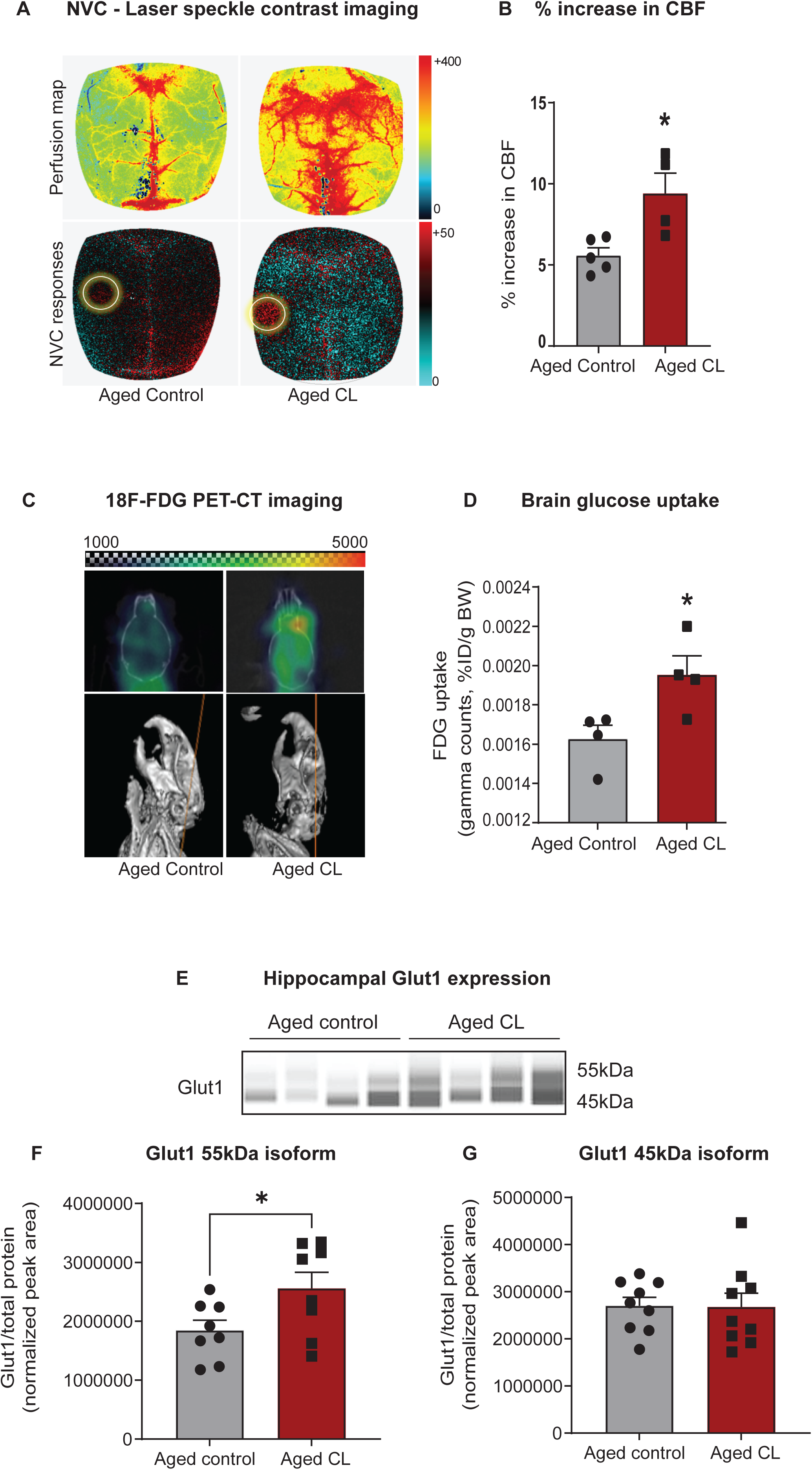
Effects of chronic β3-AR treatment on neurovascular coupling, brain uptake, and GLUT1 expression in aged mice. A. Representative pseudocolor laser speckle flowmetry maps of baseline cerebral blood flow (CBF) (upper row; shown for orientation purposes) and CBF changes in the somatosensory cortex relative to baseline during contralateral whisker stimulation (bottom row, left circle, 30 s, 5 Hz) in aged mice treated with saline (aged controls) or CL 316,243 (aged CL). The color bar represents CBF as a percent change from the baseline. Panel B shows the summary data as a % increase in CBF (*n* = 4-5 in each group). C. PET images (top) matched with the CT images (bottom) of representative aged control and CL-treated mice. Warmer colors represent higher activity in PET images. D. Quantification of FDG uptake in the brain represented as SUV (%ID/g body weight) (*n* = 4 in each group). E. Representative images of GLUT1 chemiluminescent signals in Jess capillaries created by the compass SW software for Jess analysis. F-G. Peak areas for 55 and 45kDa GLUT1 isoforms normalized for total protein in the samples. Data are mean ± S.E.M. (*n* = 8-9 in each group). \**P* < 0.05 vs. aged controls.

### Chronic β3AR stimulation attenuated BBB leakage and neuroinflammation in aged mice

Next, we evaluated the effects of β3AR stimulation on microvascular endothelial structure which critically contributes to the maintenance of BBB integrity in aged mice. To determine BBB permeability, 3kDa FITC labeled dextran was injected intraperitoneally, and the extravasation of the injected tracer was quantified in the cortex tissue lysates after perfusion. CL treatment attenuated BBB leakage in the cortex tissue of aged mice evident from the decreased permeability of the injected tracer in the brain parenchyma (Fig. 2A-B).

**Figure 2:**
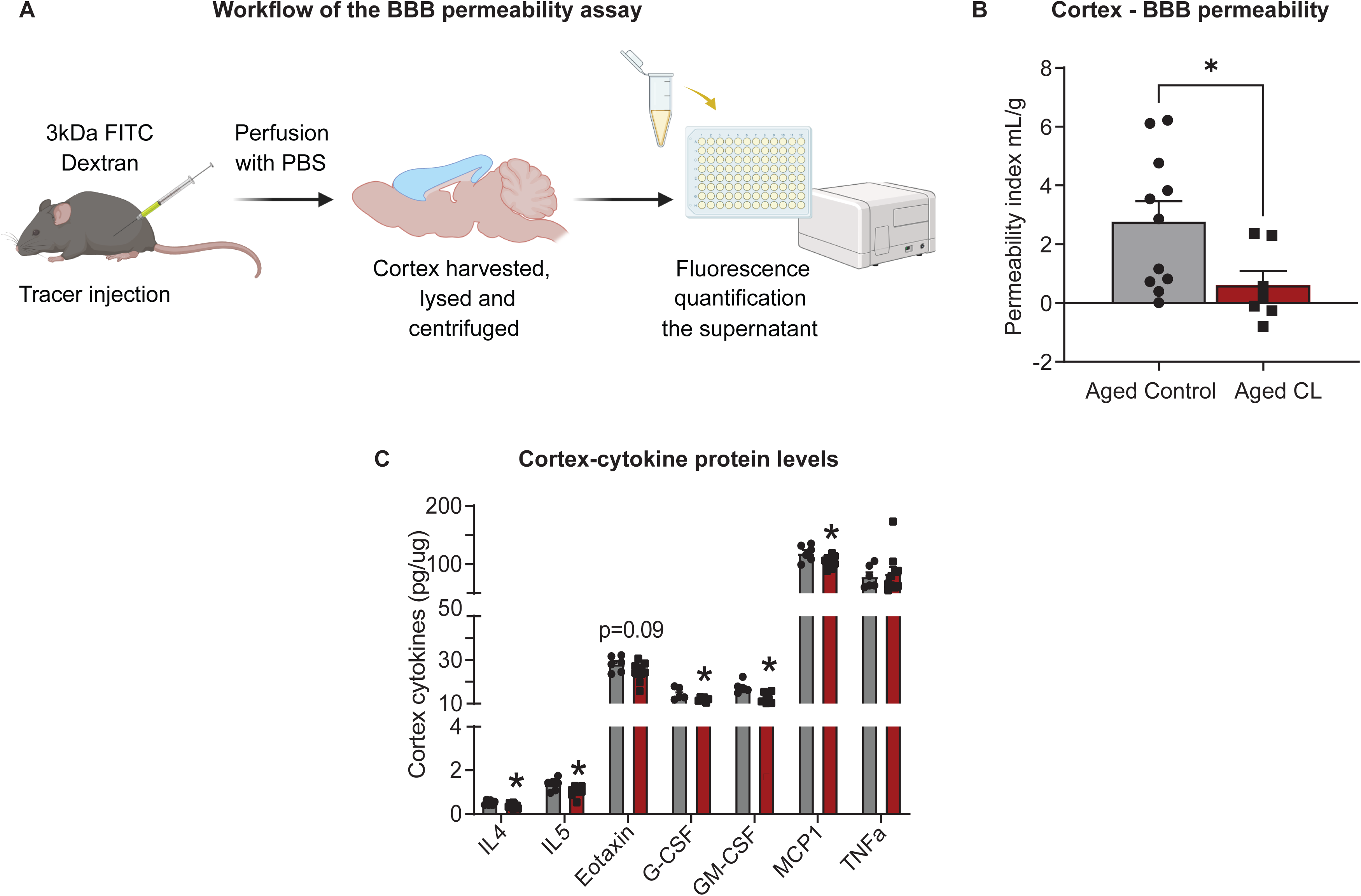
Effects of chronic β3-AR treatment on BBB permeability and inflammatory markers in the brains of aged mice. A. Workflow representing the steps in the BBB permeability assay. B. BBB permeability index calculated to assess the permeability of 3kDa FITC tracer in the cortex of aged controls and CL-treated mice (*n* = 6-11 in each group) C. Protein levels of pro-inflammatory cytokines and chemokines assessed by multiplex magnetic assay (*n* = 6-9 in each group). Data are mean ± S.E.M. \**P* < 0.05 vs. aged controls.

Intact BBB is crucial to prevent the trafficking of immune cells and other plasma proteins into the brain parenchyma. However, with aging, increased BBB permeability to blood constituents results in aberrant glial activation which ultimately leads to neuroinflammation. Hence, we wanted to assess if CL treatment mediated mitigation of BBB leakage improved inflammation in the aging brain. We assessed inflammation by measuring the protein levels of pro-inflammatory cytokines and chemokines in the cortex tissue lysates. Conforming to improved BBB function, CL treatment significantly reduced the protein levels of various pro-inflammatory mediators like IL5, Eotaxin, GCSF, GM-CSF, and MCP1 respectively (Fig. 2B). These findings indicate that CL treatment significantly improved microvascular structure and function in aged mice.

### Chronic β3AR stimulation improved spatial learning and memory in aged mice

To examine the effects of thermogenic stimulation on cognitive performance, especially spatial learning, and memory, we performed radial arm water maze and y-maze tests in aged mice after 6 weeks of CL treatment. First, we quantified the combined number of errors calculated across all the trials between the control and CL-treated aged mice. CL treatment significantly reduced the number of errors before reaching the target when compared to controls (Fig. 3B). During the learning trials, we also observed that the mice from both groups progressively took less time to find the target suggestive of task learning (Fig. 3C). On the last learning and probe trial, CL-treated aged mice took significantly less time to reach the target indicative of improved learning plasticity and memory when compared to controls. Further, a similar trend for improved relearning was also observed during the reversal trial in aged mice with CL treatment, however, they did not attain statistical significance. We did not observe any changes in swimming speed (data not shown) indicating that modulation of motor function did not contribute to the improved cognitive performance in CL-treated aged mice. Similar results supporting the cognitive benefits of thermogenic stimulation were also observed in the Y-maze test. The number of novel arm entries was significantly increased in CL-treated aged mice when compared to controls (Fig. 3C), suggesting improved spatial reference memory in response to thermogenic stimulation.

**Figure 3:**
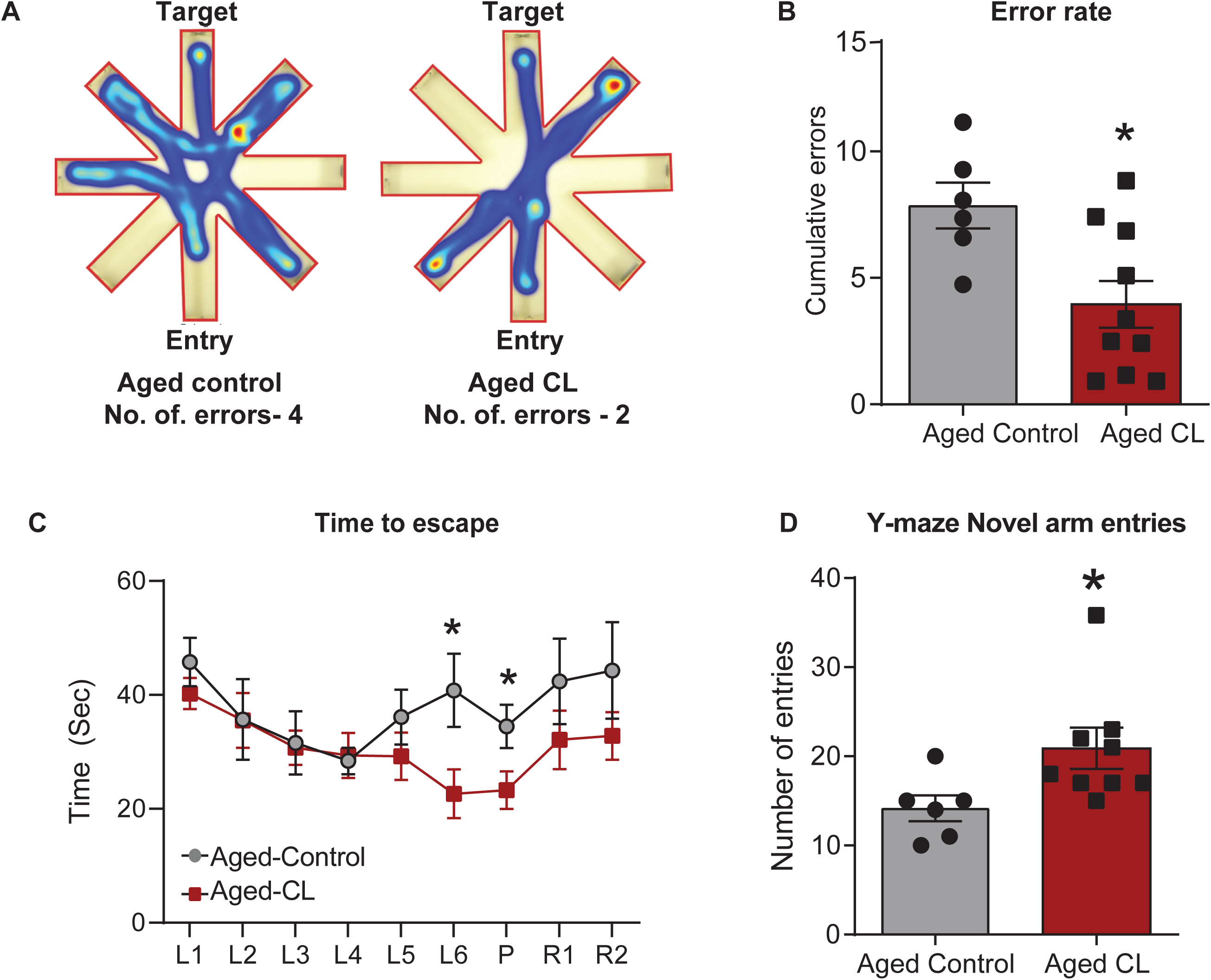
Effects of chronic β3-AR treatment on cognitive performance in aged mice. A. Radial arm water maze-Heatmap showing an animal from each group that was chosen at random and the amount of time they spent in different arms and also the traces indicating the path the mice took to reach the target. Please take note that the aged controls took longer paths and made more errors in finding the target platform when compared to the CL-treated mice. B. Cumulative errors calculated during the learning, probe, and reversal trials C. Time to escape calculated during each of the trials and D. Number of novel arm entries calculated during the Y-maze test. n=6-11 per group. Data are mean ± S.E.M. \**P* < 0.05 vs. aged controls.

## Discussion

Age-related metabolic diseases share strong pathogenic links with cerebromicrovascular dysfunction and cognitive impairment^52^. Supporting this idea, several epidemiological studies have demonstrated a causal association between metabolic syndrome in mid-life with decreased cerebral blood flow and cognitive decline later in life^39–42^. Adipose tissue dysfunction significantly contributes to the pathogenesis of metabolic disorders with aging through impaired glucose and lipid metabolism, altered adipokine secretion, increased secretion of pro-inflammatory mediators, and ectopic lipid deposition. Hence, interventions that improve adipose function and in turn peripheral metabolism might also confer protective effects on cerebral microvasculature and cognitive functions in aging. To test this, we chose thermogenic stimulation using β3AR agonists, a method previously well-established to improve adipose and systemic metabolism in both rodents and humans^53 54 55 56^. Although the effects of β3AR agonists on systemic metabolism have only been well-characterized in young animals, our studies showed that it also effectively improved multiple metabolic parameters including adiposity, glucose metabolism, insulin sensitivity, circulating adiponectin, and ectopic lipid deposition^43^. More importantly, these systemic improvements were associated with improved microvascular endothelial function and cognition in aged mice, indicating that the beneficial effects of thermogenesis extend beyond metabolic tissues. Our results are in line with previous studies which also show that CL treatment improved brown adipose tissue thermogenesis and cognition in a triple transgenic mouse model of AD (3xTg-AD)^57^ and chicks^58^, albeit the mechanisms remain uncharacterized.

We posit that the mechanisms underlying the cognitive benefits of CL treatment in aging are multifactorial. First, central insulin sensitivity likely mirrored the improvements in peripheral insulin sensitivity resulting in increased endothelial GLUT1 expression and brain glucose uptake in aged mice following CL treatment. GLUT1 is highly expressed in the brain microvascular endothelial cells^59^, where it regulates BBB integrity and cerebral blood flow responses^60^ in addition to supporting metabolic needs. Endothelial GLUT1 deficiency has been linked to impaired cerebral blood flow, BBB breakdown, and cognitive impairment in AD mice models^60^. Further, reduced GLUT1 expression anticipates the onset of microvascular dysfunction and clinical manifestations in mild cognitive impairment (MCI) and AD patients^61,62^, suggesting a pathogenic role for impaired endothelial glucose uptake in age-related vascular cognitive impairment. Based on these findings, it is highly likely that increased endothelial glucose uptake via GLUT1 (55kDa isoform) contributed to the restoration of BBB integrity leading to reduced neuroinflammation and improved learning and memory in aged mice. In addition, studies have also pointed to a role for GLUT1 in eNOS-mediated endothelial relaxation^63^ and hence GLUT1 potentially also contributes to improved neurovascular coupling observed in CL-treated aged mice. However, the mechanistic contribution of GLUT1 deficiency to neurovascular uncoupling in aging is yet to be investigated.

Secondly, the potential role of adipose-secreted factors on cerebral microvasculature should also be considered as adipose tissue is the primary tissue target for CL. Specifically, we have observed that CL treatment significantly increased the circulating levels of adiponectin, a well-known vasoprotective adipokine with insulin-sensitizing^64^ and anti-inflammatory properties^64–68^. Adiponectin has been shown to protect endothelial cells against high glucose and oxidized LDL-induced oxidative stress^69,70^, increase the production of NO by activating AMPK-eNOS signaling^11,79, 71^, and maintain capillarity and microvascular blood flow^72^. Adiponectin was also reported to inhibit atherogenesis^67^ and to modulate inflammatory processes in cerebromicrovascular endothelial cells^73^. Further, studies have also established a critical role for adiponectin in the anti-aging vascular effects of caloric restriction^74,75^. Given that adiponectin receptors (primarily AdipoR1) are expressed in brain microvascular endothelial cells^76^, it raises the possibility that adiponectin could directly influence endothelial outcomes in CL-treated aged mice. It should also be noted that adipose tissue secretome is not just limited to adipokines but includes a wide repertoire of molecules such as bioactive lipids, peptides, and extracellular vesicles. Future studies should address whether CL treatment impacted these other adipose-secreted factors to modulate microvascular function and cognition in aging.

Thirdly, CL could also directly act on the brain to confer cognitive benefits in aging. The presence of β3AR mRNA has been documented in multiple brain regions^77^, albeit at much lower levels than in adipose tissue. Amibegron, another β3AR agonist, has been shown to possess anxiolytic properties in rodents through modulation of neurotrophic and apoptotic pathways in the hippocampal neurons^78^. However, unlike amibegron which is BBB permeant, CL does not cross the BBB^79^ and it is unlikely that CL directly influenced neuronal function. Alternatively, microvascular endothelial cells are indeed exposed to CL in circulation and are a potential target for its central actions. β3AR expression in the brain microvascular endothelial cells has not been defined yet, however, it is present and physiologically active in the coronary and retinal endothelial cells^80,81^. In both the heart and retinal microvessels, stimulation of β3ARs induces eNOS activation and vasodilatory responses^80,81^. Whether β3AR stimulation exerts similar actions in brain microvessels is yet to be characterized.

Irrespective of the mechanisms, our findings show that chronic β3AR agonist treatment exerts robust microvascular protective effects in aged mice, which likely conferred cognitive benefits in aged mice. While β3AR agonists are being tested in clinical studies for metabolic disorders^44,45^, it could be a valuable therapeutic strategy to repurpose them for the treatment of age-related vascular cognitive impairment.

## Acknowledgments

This work was supported by grants from the NIH (K01AG073613), American Heart Association (CDA1048544), American Federation for Aging Research Supplement award, Presbyterian Health Foundation, and OUHSC College of Medicine Alumni Association (COMAA) to PB.

## Author Contributions

Conceptualization, DN and PB; methodology and investigation, DN, SE, ST, AH, RN and VA; writing—original draft preparation, PB; review and editing, DN, MS, ST, AC, AY, VA and PB; funding acquisition, PB.

## Data Availability Statement

The data that support the findings of this study are available from the corresponding author upon reasonable request.

## Figure legends

**Suppl. Fig. 1:**
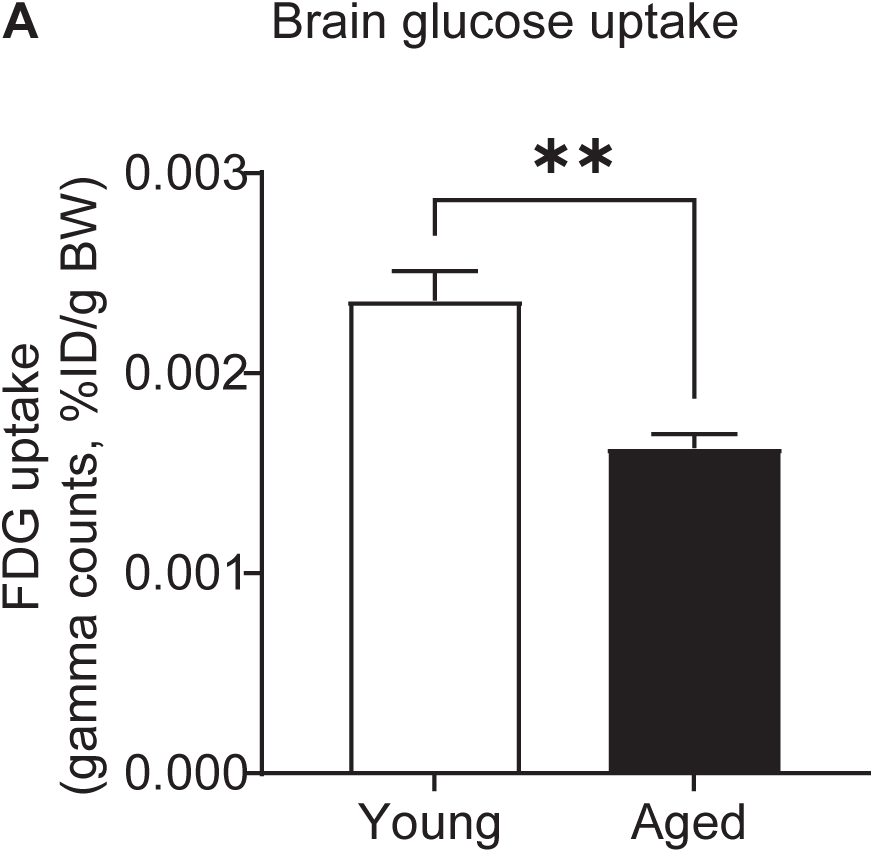
Quantification of FDG uptake in the brains of young and aged mice represented as SUV (%ID/g body weight) (*n* = 4 in each group). Data are mean ± S.E.M. \**P* < 0.05 vs. young controls.

